# Using taxon resampling to identify species with contrasting phylogenetic signals: an empirical example in Terrabacteria

**DOI:** 10.1101/369264

**Authors:** Ashley A. Superson, Doug Phelan, Allyson Dekovich, Fabia U. Battistuzzi

## Abstract

**Motivation:** The promise of higher phylogenetic stability through increasing dataset size within Tree of Life (TOL) reconstructions has not been fulfilled, especially for deep nodes. Among the many causes proposed are changes in species composition (taxon sampling) that could influence phylogenetic accuracy of the methods by altering the relative weight of the evolutionary histories of each individual species. This effect would be stronger in clades that are represented by few lineages, which is common in many Prokaryote phyla. Indeed, phyla with fewer taxa showed the most discordance among recent TOL studies. Thus, we implemented an approach to systematically test how the number of taxa and the identity of those taxa among a larger dataset affected the accuracy of phylogenetic reconstruction.

**Results:** We utilized an empirical dataset of 766 fully-sequenced proteomes for phyla within Terrabacteria as a reference for subsampled datasets that differed in both number of species and composition of species. After evaluating the backbone of trees produced as well as the internal nodes, we found that trees with fewer species were more dissimilar to the tree produced from the full dataset. Further, we found that even within scenarios consisting of the same number of taxa, the species used strongly affected phylogenetic stability. These results hold even when the tree is composed by many phyla and only one of them is being altered. Thus, the effect of taxon sampling in one group does not seem to be buffered by the presence of many other clades, making this issue relevant even to very large datasets. Our results suggest that a systematic evaluation of phylogenetic stability through taxon resampling is advisable even for very large datasets.

**Supplementary information:** Supplementary text and figures are available on the journal’s website.

## 1 Introduction

The promise of phylogenomic approaches consisted in higher phylogenetic accuracy and stability even in the presence of incomplete datasets (e.g., missing data or incomplete taxon sampling) (e.g., Felsenstein, 1985; Huelsenbeck and Rannala, 1997; Boore, 2006). This promise stems from an increased availability of sequenced genomes and faster algorithms for the analysis of thousands of species, which support the reconstruction of increasingly more inclusive Trees of Life (TOL). In an ideal case scenario, these TOL should be able to represent the full extent of the evolutionary processes of life over four billion years (Delsuc *et al.*, 2005). Phylogenomic approaches have been used widely in both Eukaryotes and Prokaryotes but it is especially in Eukaryotes that increased samplings have resolved the phylogenetic placement of groups that are taxonomically diverse and that have been phylogenetically unstable with smaller datasets (Burki *et al.*, 2016). For example, deep relationships within the phylogenies of insects (Shin *et al*, 2018) and angiosperms (Massoni, 2014) have shown increased resolution when taxon numbers are increased. However, TOL reconstructions have also shown poor phylogenetic convergence for many important nodes especially in the deepest parts of the trees, raising doubts that higher accuracy can be achieved purely by larger datasets (sites and taxa) (Shen, 2017).

Within trees of life, relationships among prokaryotic phyla are particularly unstable most likely because of compounding effects of horizontal gene transfers, difficult orthology determination, inaccurate evolutionary models, and compositional biases (Som, 2014). All of these factors are inherently linked to taxon sampling because different species will have unique gene histories and other biases (e.g., compositional biases) that will weigh differently on the overall phylogenetic signal. Thus, evaluating phylogenetic instability under different taxon re-samplings can lead to the identification of lineages or groups of lineages with contrasting phylogenetic signals. While, traditionally, analyses of taxon sampling have evaluated the effect of number of taxa (e.g., Rokas *et al.*, 2003; Gatesy *et al.*, 2007; Heath *et al.*, 2008), the estimation of the relative effect that each species has, when keeping the same number of species, is equally important. This approach leads to more focused hypothesis-testing when searching for causes of the conflicting signals because it allows for applied analyses of sequence properties (e.g., compositional biases, rate of evolution) to those species with diverging signals (Gatesy *et al.*, 2007). These fewer and more streamlined tests can then be applied easily to more extensive datasets enabling comprehensive evaluations of the robustness of the conclusions based on the obtained tree.

As an example of this proposed approach we focus on recent studies by Lang *et al.* (2013), Rinke *et al.* (2013) and Hug *et al.* (2016), each of which utilized large datasets of microbial species to reconstruct a TOL. These TOL show overall agreement for the monophyly of domains and phyla but disagree in their relative relationships despite broad similarities in dataset sizes and methodologies. For example, all three maximum-likelihood phylogenies confirm the strong associations among phyla of the Planctomycetes, Verrucomicrobia, and Chlamydiae, and show a well-known resolved monophyly for this superphylum (Wagner *et al.*, 2006). However, clades formed by phyla in the Terrabacteria superphylum, are inconsistent (Fig. 1). A possible explanation for these results is that the differences in taxon sampling among these studies (e.g., Hug *et al*. (2016) used 6 Deinococcus-Thermus while Lang *et al*. (2013) used 25) could be a leading cause of their inconsistency with each other. These effects are likely to be amplified in those phyla represented by few lineages where the weight of a single species is proportionally larger than what it would be in a group represented by hundreds of species. Indeed, a closer analysis of the phylogenetic disagreements in these three studies shows that phyla represented by few species (e.g., Deinococcus-Thermus) have the most variable phylogenetic placement, raising the possibility that the incongruence seen across reconstructions could be directly correlated with taxon sampling and, indirectly, with the evolutionary histories of the species used to represent a clade. Working under the simple hypothesis that the instability of the Terrabacteria group and subgroups is caused by the number of species used and/or the identity of the species chosen to represent each group, we created alternative scenarios with increasingly fewer taxa for one clade at a time to investigate the stability of the phylogeny in light of different species compositions. In general, if taxon sampling (and the correlated lineage-specific properties) has no influence on phylogenetic reconstruction we would expect all altered datasets to recover the same topology as the complete, unaltered dataset.

**Figure 1.**
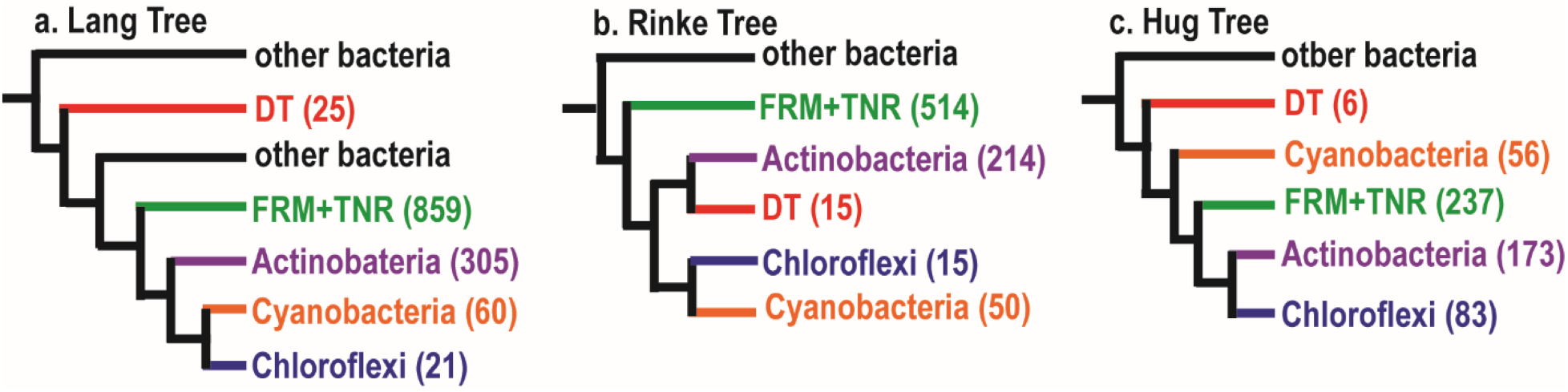
Published Maximum Likelihood phylogenies showing backbone of Terrabacteria phyla. In parenthesis next to each phylum is shown the number of species used for each dataset and group. **a.** Tree modified from Lang et al. (2013). **b.** Tree modified from Rinke et al. (2013). **c.** Tree modified from Hug et al. (2016).

To test this, we created a phylogenetic pipeline that removed a user-defined number of species within a single phylum from a concatenated alignment of orthologous genes, while keeping all other phyla intact (Fig. 2). The pipeline retains the data of intermediate steps for subsequent analyses to evaluate other variables (e.g., composition bias of the alignment) and parameters (e.g., alignment length) that are inherently altered when taxon sampling is changed. We found that both taxon sampling and species identity impact the overall congruence in phylogenetic reconstruction. Additionally, we found that neither alignment length nor the compositional biases specific to each set of representatives were the primary causes of the detected incongruences. Based on these results, we also identified a minimum of 15 species within Deinococcus-Thermus, Chloroflexi, and Actinobacteria that produce contrasting phylogenetic signals compared to the rest of the species.

**Figure 2.**
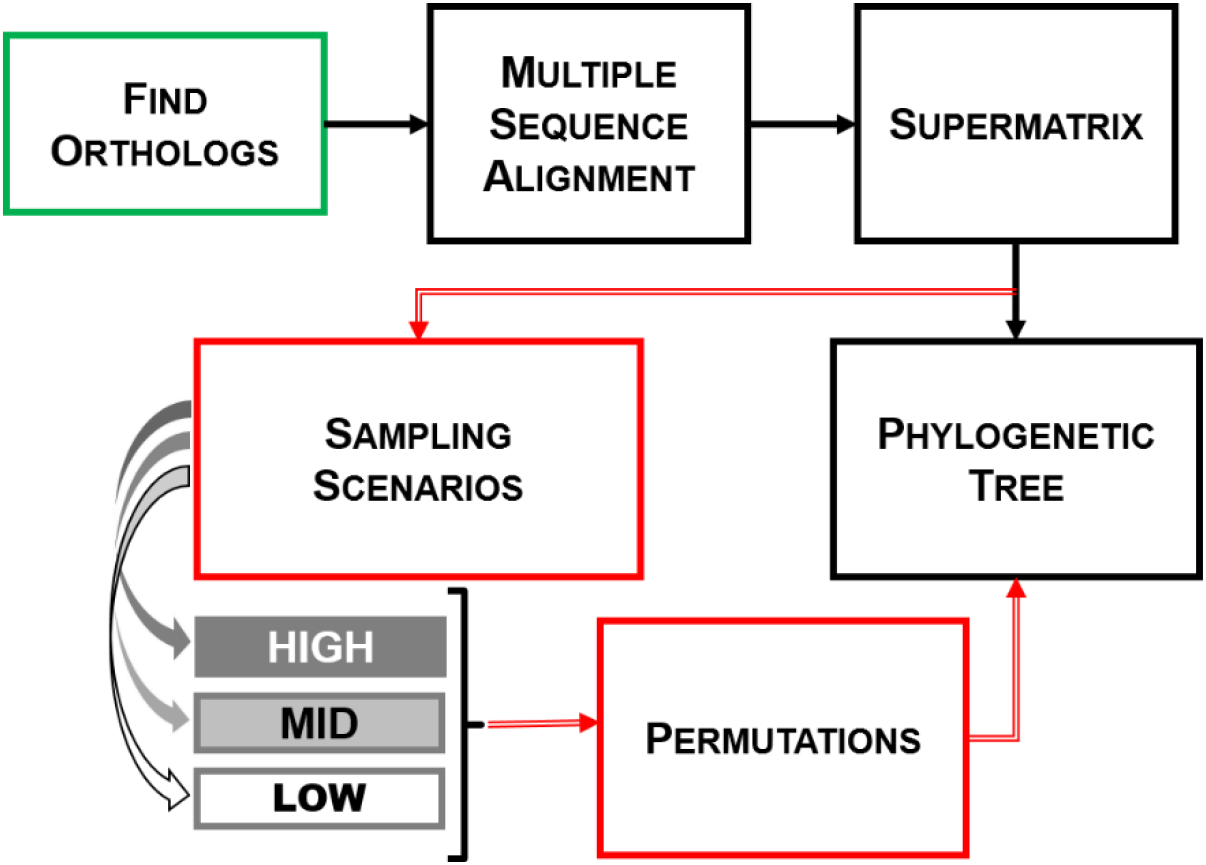
Phylogenetic pipeline to produce taxon resamplings. Standard reconstruction steps are indicated through solid black arrows. To test hypotheses about the effect of taxon sampling within this process, we implemented the steps shown in red that create resamplings with different species numbers (“sampling scenarios”) and different lineages with the same number of species (“permutations”). Sampling and permutations were implemented either after the alignment step or from the beginning by reassessing orthologs (green box).

## 2 Methods

### 2.1 Dataset and phylogenetic pipeline

The initial dataset was composed of fully sequenced proteomes for six phyla within the Terrabacteria superphylum available from the National Center for Biotechnology Information as of 7 March, 2016. Species with multiple strain representatives were manually filtered to include only the strain that had the largest genome, resulting in a total of 766 species (we refer to this dataset as FULL). We analyzed this dataset to explore the effect on phylogenetic reconstruction of two variables: (i) the number of species present in a given phylum and (ii) the identity of these species. We automated a phylogenetic pipeline that implemented standard phylogenetic reconstruction approaches (i.e., orthology determination, alignment, tree building) on different sampling scenarios (HIGH, MID, LOW; see below) that modified the taxon sampling within a desired phylum (Fig. 2). Additionally, each scenario was iterated exhaustively in the two small phyla analyzed and heuristically in a large phylum to explore the effect that each individual species or group of species might have on the reconstructed phylogeny (Fig. 2: red arrows). After evaluating available options, software and parameters selections for the pipeline were systematically optimized based on computational demands and overall performance (Supplementary Data).

ProteinOrtho v5.16b (Lechner *et al.*, 2011) was used to identify homologous genes based on results of bidirectional blastp scores at 0 connectivity. We used an in-house script to filter the homologous groups and identify orthologous groups based on a varying threshold of species represented in each ortholog. We tested multiple thresholds (100%, 99%, 97%, 95%, 90%, and 80%) and found that the 90% threshold optimized the number of orthologs (30 orthologs) and amount of missing data (< 2%) (Supplementary Data). Each ortholog was aligned using MUSCLE (Edgar, 2004) under default parameters. The alignments were filtered to exclude sites with more than 25% gaps. After concatenating each individually aligned ortholog into a supermatrix, we obtained an alignment consisting of 6,227 sites.

The Terrabacteria dataset provided a good representation of the bias seen in the availability of sequenced genomes. Within the Terrabacteria, Actinobacteria had 265 fully sequenced species, Firmicutes (which, for the purpose of this study, include also Tenericutes) 394, Deinococcus-Thermus 20, Chloroflexi 14, and Cyanobacteria 73. We implemented different sampling scenarios to the alignment by removing a set number of taxa for a chosen phylum (e.g., Deinococcus-Thermus) without altering the number of species present for other phyla. Among these phyla, we chose the two smallest ones (DT and CHF) to test changes in their phylogenetic position within each scenario and Actinobacteria as a control group with a large number of species.

Due to the heavy computational requirements necessary to identify the best substitution model, we applied ProtTest v3.4 (Darriba *et al.*, 2011) to 15 datasets with 100 species, which were randomly produced from the original concatenation. In all cases the most complex model (LG) was found to be the best fit. We then implemented this model in FastTree (Price *et al.*, 2010) to estimate ML phylogenies for all scenarios and permutations.

### 2.2 Analytical framework

Trees were arbitrarily rooted using the Cyanobacteria phylum and each tree was analyzed at the backbone and all internal nodes levels.

At the backbone level, we collapsed taxa that are classified in the same phylum to easily evaluate the relationship among phyla in the estimated tree. In the case of the backbone analyses, all phyla were collapsed so that each tree is composed of 6 branches that represent the 5 phyla. The sixth branch represents an exception made for two species, *Coprothermobacter proteolyticus* (CTP) and *Thermodesulfobium narugense* (TSN) that despite being classified as Firmicutes, consistently clustered outside of this phylum and thus were collapsed independently of their phylum.

At the internal nodes level, only the permutated phylum (DT, CHF, or ACT) was collapsed leaving the other phyla represented by all their species. To evaluate the similarity among trees produced from our pipeline we used Robinson-Foulds (RF) distances obtained with IQ-TREE (Nguyen *et al.*, 2014). We treated all the trees produced under a given permutation scenario (i.e., DT, CHF, or ACT) as a set and standardized the results using the maximum RF value possible based on number of nodes (nRF). This allowed us to directly compare the results among permutation scenarios. Additionally, because each sampling and permutation scenario has a different number of trees that need to be compared, we weighted the standardized RF scores by the number of trees in each category (nRF*) (Supplementary Data).

### 2.3 Taxon sampling in small groups (DT and CHF)

The LOW sampling scenario is obtained by removing all species, except for one, from the dataset for either the DT or the CHF phylum resulting in 747 total species in the DT permutations and 753 species in the CHF permutations. HIGH taxon sampling was represented by the inverse, where 19 of 20 Deinococcus-Thermus and 13 of 14 Chloroflexi species were kept resulting in trees consisting of 765 species. The MID level sampling scenario kept groups of species that were monophyletic at the genus level and had more than one representative. Deinococcus-Thermus had two groups, the genus Deinococcus represented by 10 species and the genus Thermus represented by 6 species (resulting in trees of 756 species and 752 species, respectively). Chloroflexi had four genera groups: Chloroflexus with 3 species and three genera with 2 species each: Dehalococcoides, Dehalogenimonas, and Roseiflexus (resulting in trees of 753 species and 754 species, respectively).

Within each sampling scenario, all available combinations of species compositions were implemented. For each HIGH and LOW scenario, each species within a given phylum was either kept or removed resulting in 20 permutations for Deinococcus-Thermus and 14 permutations for Chloroflexi, per scenario. Each permutation resulted in a tree that was then compared to the others within that scenario in order to see if the identity of the species isolated had an effect on the topology of the tree. The number of permutation trees under the MID scenario depended on the number of groups identified (two for DT, four for CHF).

### 2.4 Taxon sampling in large groups (ACT)

LOW taxon sampling was represented by keeping 1% (3 species) of the 265 species of Actinobacteria in the alignment and removing the remaining 262 species leaving 504 total species in resulting trees. For HIGH taxon sampling, we removed 10% of species from the Actinobacteria phylum which resulted in an alignment of 740 species. There were 20 permutations generated for each LOW and HIGH sampling scenario using randomly selected species. MID level sampling considered classes that contained more than 10 species and whose nodes were monophyletic in the FULL phylogeny. There were 5 permutations generated by keeping only the species within respective classes and removing all other species in the Actinobacteria phylum. The following classes were kept: Streptomycetales (32 species), Pseudonocardiales (17 species), Propionibacteriales (10 species), Corynebacteriales (86 species), and Bifidobacteriales (19 species) (species names for each scenario tested are available in the Supplementary Data).

### 2.5 Additional Considerations

The phylogenetic comparisons we perform are within a relative framework that is delimited by a tree with the full set of taxa and trees with reduced taxa sets. Although this provides consistency in the variables tested (i.e., taxon sampled), it also implies that in altering the sampling the orthology assessment, and related difference in alignment length, does not influence topology. To evaluate this assumption, we extended the pipeline to (i) re-estimate orthologous groups from each permutation dataset at the first step of the pipeline (Fig. 2, green box) as opposed to permutating species from the supermatrix and (ii) bootstrap (50 times) the length of resulting alignments to the same number of sites as the full dataset (Supplementary Data).

## 3 Results

We considered the tree produced by the FULL dataset (766 species) as the reference to compare the results of all subsequent analyses with altered species compositions. The relative relationships among taxa were established after arbitrarily rooting the tree with Cyanobacteria. Analyses of trees for both small (Deinococcus-Thermus, DT; 20 species, and Chloroflexi, CHF; 14 species) and large (Actinobacteria, ACT; 265 species) groups were considered (i) per sampling scenario, to test the influence of taxon sampling (LOW, MID, and HIGH), and (ii) per permutation scenario, to consider the effects of unique species composition and evolutionary history given by each individual lineage. Sampling scenarios tested included three levels of species representation: a HIGH scenario in which only one (or up to 10%) species was removed, a LOW level in which all species but one (or up to 99%) were removed, and a MID level in which groups of monophyletic species were removed (Methods). For each scenario, multiple permutations were performed to exhaustively or heuristically analyze a large number of possible combinations. To summarize general trends, we evaluated relationships at three levels: the closest relative of Deinococcus-Thermus, the backbone of the tree, and all internal nodes for each of the trees produced.

The backbone topology produced when all 766 species (FULL) were used is shown as tree A in figure 3, where species within a monophyletic phylum collapsed within each branch. Collapsing the monophyletic groups at the phylum level from our dataset leaves 5 phyla as the backbone, assuming Firmicutes and Tenericutes as a single branch. *Coprothermobacter proteolyticus* (CTP) and *Thermodesulfobium narugense* (TSN) were consistently not clustering according to their classification (Firmicutes) but forming a monophyletic group of their own, which we counted as a separate, sixth clade (Fig. 3a, light green branches). Because of this unusual placement, we repeated some of the analyses without these two species (below; Fig. 3b). Excluding Cyanobacteria that we used as outgroup, there are 15 possible different unrooted topologies with 5 phyla (105 with 6 phyla) (Felsenstein, 1978), but, throughout our analyses, we recovered only seven topologies (Fig. 3a).

**Figure 3.**
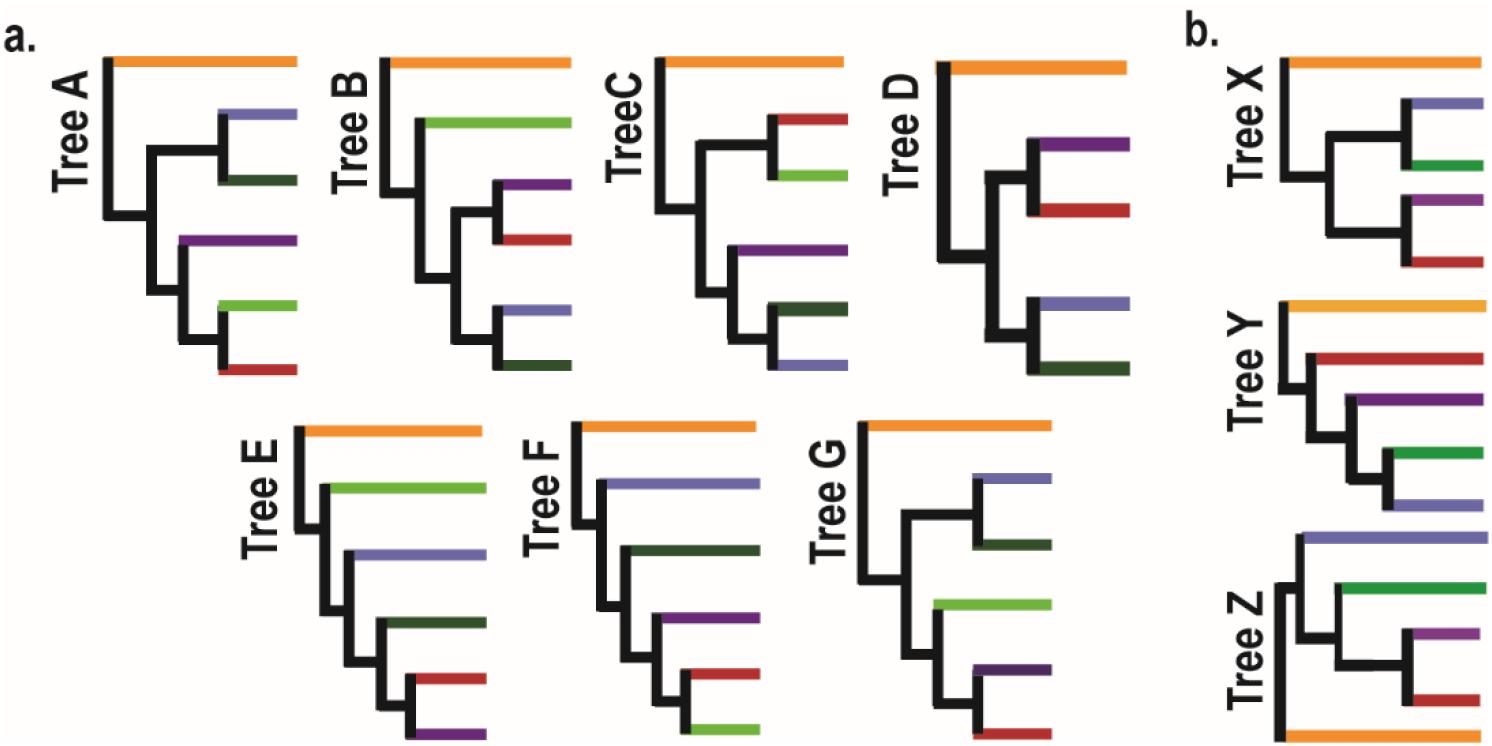
Backbone topologies. Each branch represents species within a given phylum of Terrabacteria: Cyanobacteria (orange), Chloroflexi (blue), Actinobacteria (purple), Deinococcus-Thermus (red), and Firmicutes/Ternericutes (green). **a.** Trees A-G show the different backbone topologies of all trees produced. Within these trees Firmicutes are paraphyletic; the two species clustering outside Firmicutes/Tenericutes (dark green) are *Coprothermobacter proteolyticus* (CTP) & *Thermodesulfobium narugense* (TSN) (light green). **b.** Trees X,Y,Z show the reduced set of topologies represented without the CTP/TSN branch.

### 3.1 Comparisons of closest relative of Deinococcus-Thermus

Initially, we measured tree similarities based on the identity of the closest relative of Deinococcus-Thermus (DT). The topology produced from the FULL dataset identified the closest relative to DT as the CTP/TSN cluster (Fig. 3: tree A). Based on the 119 trees generated by the combination of sampling and permutation scenarios for both small and large groups, we found 40% of trees resulted in Actinobacteria as the closest relative to DT, while the other 60% clustered DT with CTP/TSN (Fig. 4a: black bars). When we considered these results per sampling scenario, we see that this signal is largely driven by the frequencies in HIGH sampling (i.e., > 96% of trees show CTP/TSN as the closest relative of DT) (Fig. 4a: dark grey bars). In MID and LOW sampling, the phylogenetic signal flips to approximately 70% of trees resulting in Actinobacteria as the closest relative to DT (Fig. 4a: light grey and empty bars).

**Figure 4.**
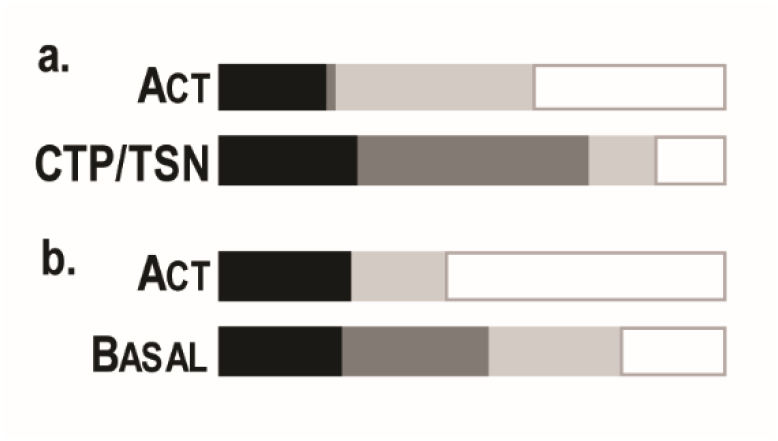
Deinococcus-Thermus (DT) closest relative frequencies. Frequencies (0 – 100% for combined bars) are represented per sampling scenario for 119 trees produced: FULL (black), HIGH (dark grey), MID (light grey), LOW (empty). **a.** Results of trees produced with *Coprothermobacter proteolyticus* (CTP) & *Thermodesulfobium narugense* (TSN) where DT either had Actinobacteria (ACT) or CTP/TSN as its closest relative. **b.** For trees produced if CTP/TSN branch was not included, either ACT is closest relative to DT or DT is basal to the rest of the ingroup phyla.

When we analyzed the results of the permutations within each sampling scenario, we saw similar trends across HIGH sampling for all permutations with more than 90% of the permutation trees showing CTP/TSN as the closest relative to DT. The MID sampling scenario in Actinobacteria also showed that the trees produced by the permutations are consistently in agreement with the FULL dataset; however, in the small groups analyses (DT and CHF) the permutations result in a signal that is evenly split between CTP/TSN and ACT. Within LOW permutations, instead, the majority (> 90%) of the permutations on Chloroflexi and Actinobacteria showed Actinobacteria as the closest relative to DT, while the Deinococcus-Thermus permutations showed 70% of the permutations with CTP/TSN as their closest relative (Table 1). Thus, the closest relative of DT varies between CTP/TSN and ACT across sampling and permutation scenarios with lower taxon sampling favoring a clustering different from the high taxon sampling results.

**Table 1.**
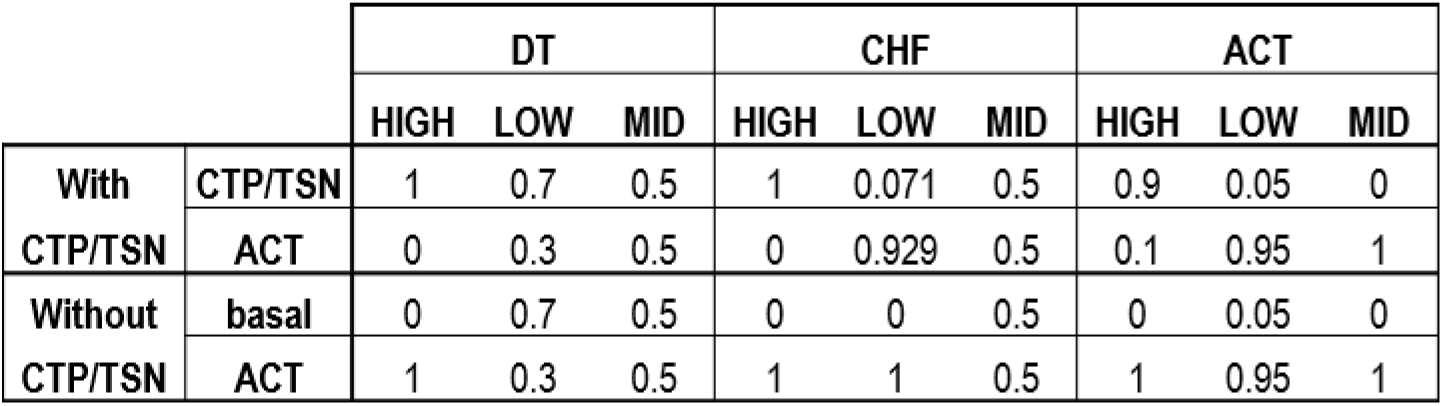
Deinococcus-Thermus (DT) closest relative frequencies. Frequencies are represented per permutation scenario (DT, Chloroflexi (CHF), and Actinobacteria (ACT)) for 118 trees produced. For trees produced with *Coprothermobacter proteolyticus* (CTP) & *Thermodesulfobium narugense* (TSN), DT either had Actinobacteria (ACT) or CTP/TSN as its closest relative. If CTP/TSN branch were not present, DT either would have ACT as its closest relative or move basal to the rest of the ingroup.

### 3.2 Comparisons of backbone nodes

We then considered tree similarities based on the relationship of all backbone groups. Across all sampling scenarios, less than 50% of the trees produced the same backbone topology as the FULL tree (Fig. 3a: tree A). As seen above, this result is largely driven by the HIGH sampling scenario, as tree A is not recovered by any other sampling scenario (Fig. 5a: dark grey bars). In HIGH sampling, the permutations of small groups had no effect on the backbone recovered (i.e., taxon composition had no influence); however, taxon composition did have an influence on 10% of the trees recovered in our large group (Fig. 5b). As expected, the most discord is seen in LOW sampling where the chosen phylum is represented by the lowest number of species (Fig. 5a, empty bars). The MID sampling scenario also shows discordance, but the most common signal obtained is not congruent with the most common signal from either HIGH or LOW sampling (e.g., tree G in MID and trees B & C in LOW; Fig. 5a). Therefore, within both LOW and MID sampling, taxon composition strongly influenced the backbone recovered and revealed conflicting signals irrespective of the size of the group being altered (small or large).

**Figure 5.**
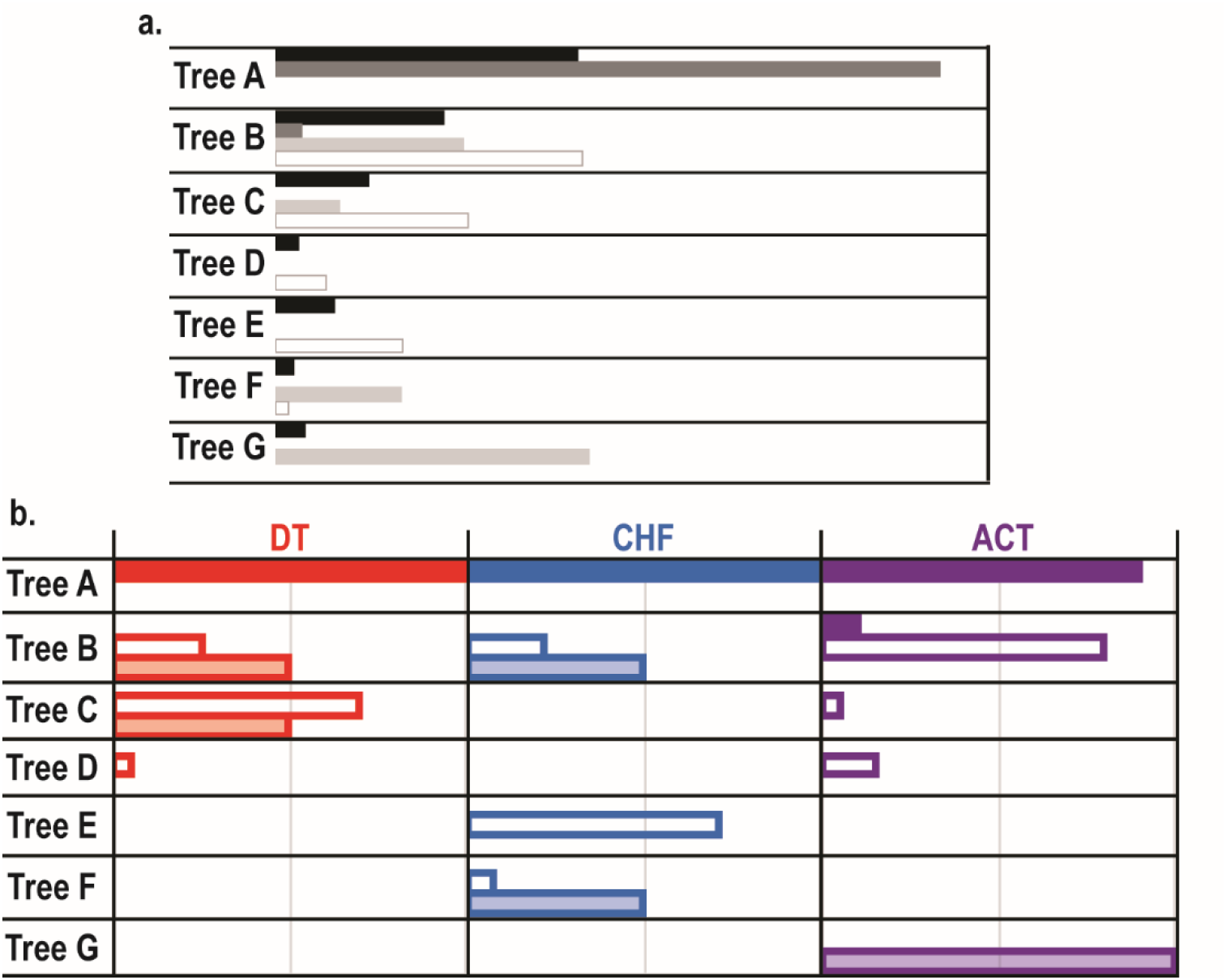
Backbone frequency results. Trees A-G as shown in figure 3. **a.** Frequencies (0 – 100% for each bar) are represented per sampling scenario for 119 trees produced: ALL (black), HIGH (dark grey), MID (light grey), LOW (empty). **b.** Frequencies are represented per permutation scenario (Deinococcus-Thermus (DT), Chloroflexi (CHF), and Actinobacteria (ACT)) for 118 trees produced for each sampling scenario (HIGH (solid), MID (shaded), and LOW (empty)).

Because all backbone analyses were conducted on alterations of the original alignment (FULL: 6227 sites; Methods and Supporting Data), we evaluated the effects that new alignments obtained from different sets of orthologs might have on phylogenetic reconstructions. We focused on the Deinococcus-Thermus (DT) phylum under LOW sampling because it is the one that produced the least stable results and evaluated (i) changes in orthologous groups and (ii) changes in the composition of alignment sites. Removal of 19 out of 20 DT species prior to the orthology assessment step, resulted in the addition of one ortholog for 19 of the 20 permutations, producing alignments that were on average 10% longer. Whether we used these longer alignments or reduced their length by bootstrapping the sites to the same length of the FULL alignment (Methods), we found that the backbone topology recovered remained the same in approximately 55% of the trees (Supplementary Data).

Because CTP/TSN clustered outside their current classification (Firmicutes) and created an additional branch in the backbone tree, we evaluated our patterns in the absence of these two species. Simply eliminating the branch containing CTP and TSN from our backbone trees (tree A-G), showed that the FULL tree, as well as most of the trees produced from the permutations across all sampling scenarios, recovers ACT as the closest relative to DT (> 85%) while the others place DT basal to the ingroup. Despite this reduction in variance seen among the backbone (from seven trees to three; Fig. 3b), the overall patterns of discordance among sampling and permutation scenarios observed with and without CTP/TSN remain (Fig. 4b). We repeated our analyses eliminating both these species, and each individually, from our original 766 species dataset, and we found that these species, together and individually, alter the topologies recovered in unique ways (Supplementary Data).

### 3.3 Comparisons of internal nodes

We extended our evaluation to whole trees using the Robinson-Foulds (RF) distance metric as a measure of tree similarity (Robinson and Foulds, 1981). This metric compares two trees with equal taxa and scores the trees according to the number of nodes that are different between them (i.e., a greater RF distance between trees is obtained for more dissimilar trees). When we applied this strategy to a set of trees, RF scores were calculated for the comparison of each tree to another, ultimately generating a RF matrix for a given set (Methods). Because of the different numbers of taxa and trees generated by the sampling and permutation scenarios, we normalized (nRF) and weighted the RF scores (nRF*) according to the size of each dataset.

Based on these standardized scores we found that, across all sampling scenarios, permutations of Actinobacteria (large groups) resulted in three-fold higher dissimilarity among internal nodes than in small groups (Actinobacteria: 15.29, Deinococcus-Thermus: 4.76, Chloroflexi: 4.36). When we compared the dissimilarity among trees obtained by each sampling scenario to the FULL, we observed that trees produced under LOW sampling resulted in the most discord for all three of our permutation scenarios, as already seen in the backbone analysis (Fig. 6). However, the discord seen within Chloroflexi is approximately 3 times lower than either Deinoccocus-Thermus or Actinobacteria (Fig.6: CHF, blue bar). MID sampling generally produces higher RF scores than HIGH except for DT.

**Figure 6.**
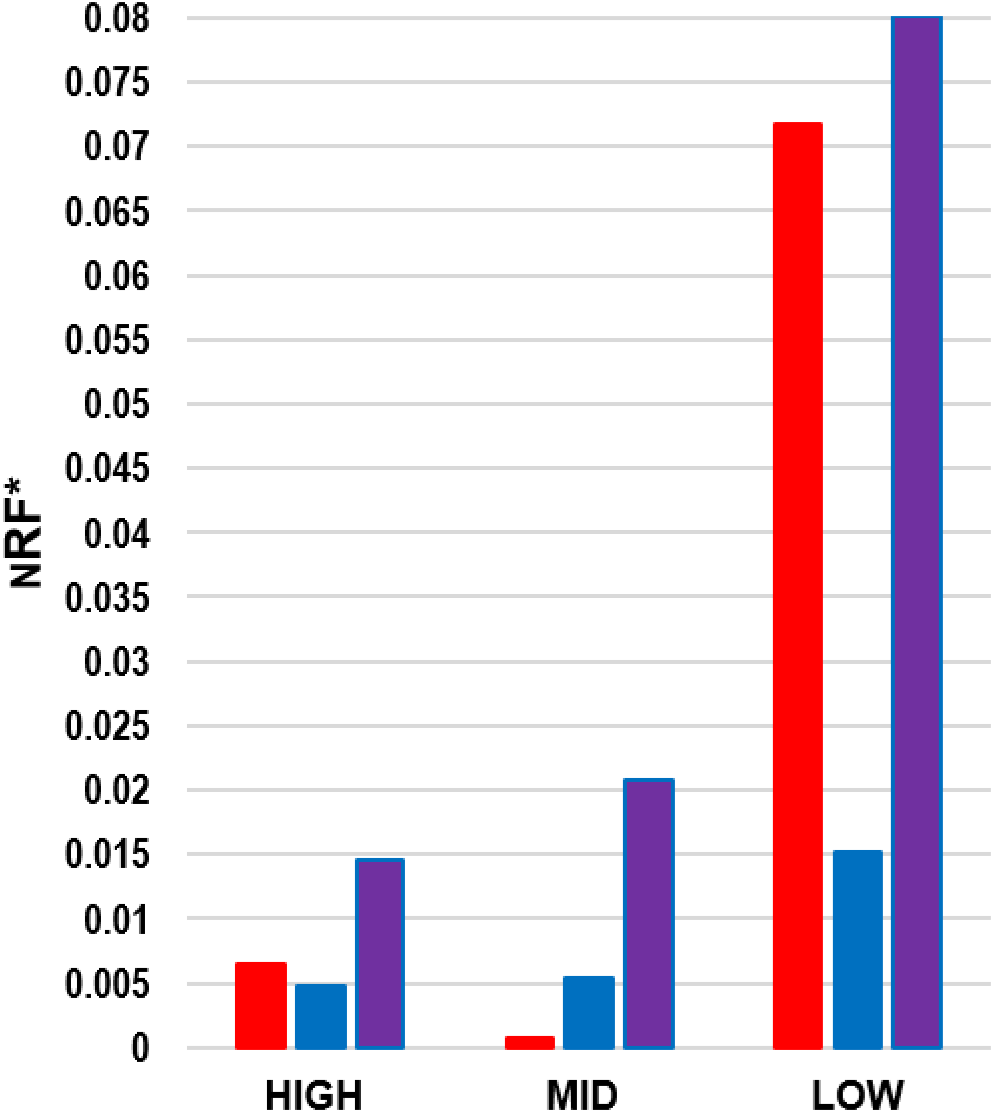
Comparison of Robinson-Foulds (RF) scores from sampling scenarios (HIGH, MID, LOW) to FULL tree. Results are shown for all three permutation scenarios ((Deinococcus-Thermus (red), Chloroflexi (blue), and Actinobacteria (purple)). RF scores were considered for the comparison of FULL tree to each permutated tree, scored were normalized (nRF) and weighted according to number of trees produced within each sampling scenario (nRF*).

Finally, through our permutation approach we identified a minimum of 15 species in the LOW sampling scenario that show contrasting phylogenetic signals compared to the remaining species. Heatmaps of tree similarity clearly show that there are distinct species that result in higher discord even when the number of taxa is the same (Fig. 7: panels a-i). Within LOW sampling, the majority of these species (9) belong to the ACT, which is not surprising given that this is our largest group analyzed. Of these, four are Corynebacteriales and one within their closely related Pseudonocaridiales, while the others are deeper within the Actinobacteria tree and include the Micrococcales (3) and Bifidobacteriales (1) (Gao and Gupta, 2012). Within DT, the species identified is *Meiothermus silvanus*. This species along with its sister lineage Thermus shares an ancestor with *Marinithermus hydrotemalis/Oceanithermus profundus* placing them towards the tips of the DT phylogeny (Ho *et al.*, 2016). Four out of the five CHF species identified belong to the Dehalococciodaceae family (includes *Dehalococcoides* and *Dehalogenimoas* genera), in which previous studies have shown unique genomic features compared to the other Chloroflexi species (Gupta *et al*, 2013). The fifth CHF species is *Anaerolinea thermophila* whose phylogenetic position varies from basal in ML trees to closer to the tips in 16S rRNA trees (Gupta *et al.*, 2013).

**Figure 7.**
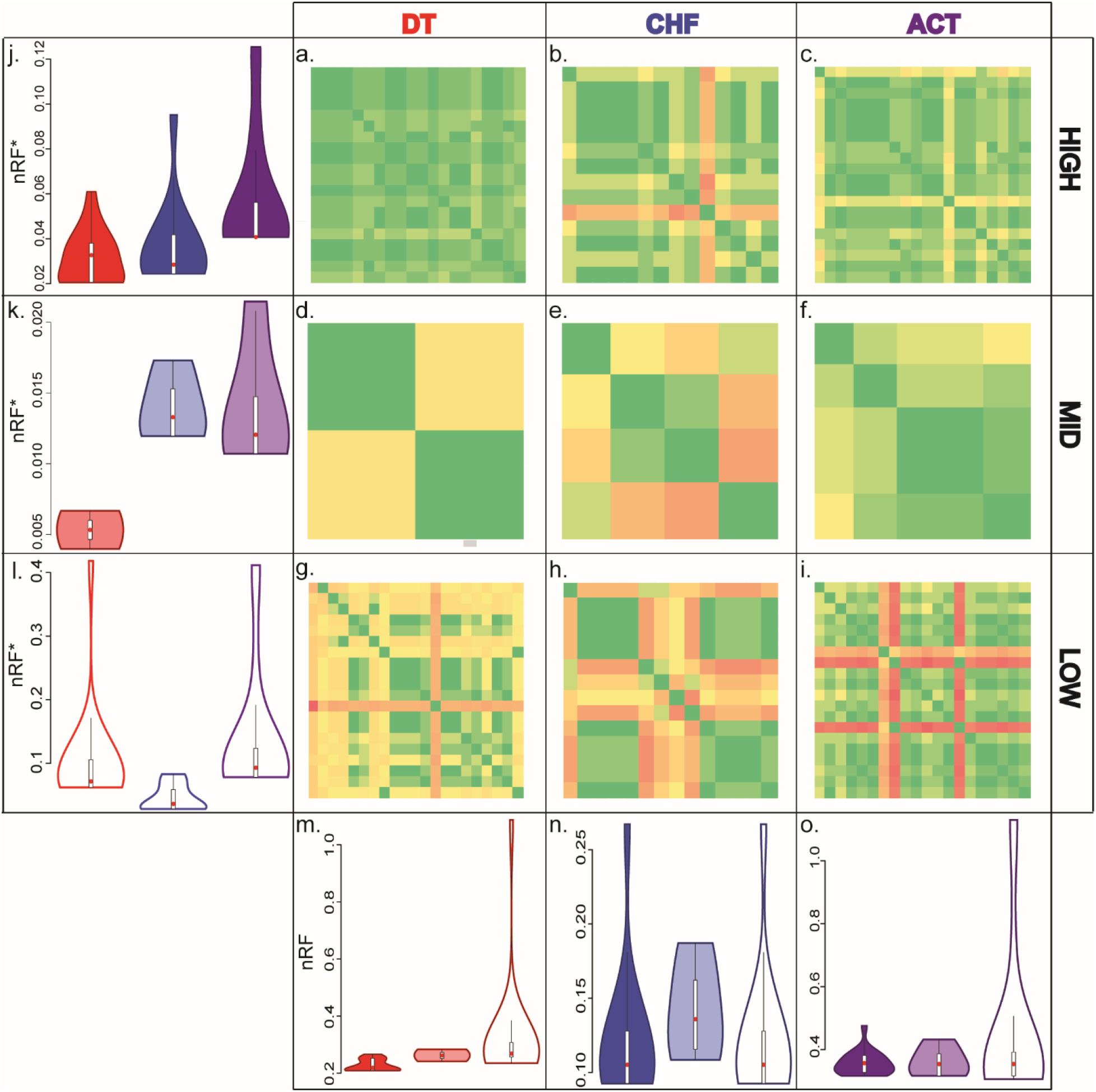
Discord among permutations and sampling scenarios. Discord among 118 trees where quantified per permutation scenario (red: Deinococcus-Thermus (DT), blue: Chloroflexi (CHF), purple: Actinobacteria (ACT)) using Robinson-Foulds (RF) metric. Scores were normalized (nRF) for each phyla permutation and weighed (nRF*) to compare across different sampling scenarios (HIGH (solid), MID (shaded), and LOW (empty)). **a-i.** Heatmaps of nRF scores corresponding to the given phyla permutations (top) and sampling scenarios (right). **j-l.** Violin plots of nRF* scores showing variance resulting from phyla permutations under a given sampling scenario (right). **m-o.** Violin plots of nRF scores showing variance resulting from altered sampling under a given permutation scenario (top).

## 4 Discussion

Recent TOL studies still show many incongruences especially in deep nodes despite the use of large-scale datasets and the complexity of the approaches applied to closely model evolutionary processes. While many studies share similar broad taxonomic sampling, details of numbers and identity of lineages used as representative(s) of each clade differ. In addition, some clades are represented by hundreds of lineages and others by only a few. This variability in taxon sampling led us to hypothesize that it may be the cause of the discrepancies in TOL reconstructions even when datasets and methods are overlapping. The idea that taxon sampling can negatively affect reconstruction efforts has been extensively tested in previous studies (Zwickl, 2002; Hedtke, 2006), primarily focusing on eukaryotes or simulated data. However, many of the phylogenetic disagreements observed in TOL studies involve the deepest nodes of Bacteria and Archaea (e.g., Lang *et al.*, 2013; Rinke *et al.*, 2013; Hug *et al.*, 2016), which can evolve using different evolutionary processes compared to eukaryotes. For example, compositional biases are more extreme in prokaryotes as is the frequency of HGT and challenges in identifying true orthologs (e.g., Jain *et al.*, 1999; Daubin *et al.*, 2002; Som 2014). Ultimately, the effect of taxon sampling is interconnected with these underlying evolutionary processes because each species used to represent a clade will inevitably carry its own genomic properties and unique evolutionary history (Gori *et al.*, 2016). While this can make it difficult to isolate the effect of each individual genomic characteristic, it allows the use of simple taxon re-sampling procedures to identify those lineages that show discordant phylogenetic signals and flag them for further analyses to determine the ultimate cause of these signals.

### 4.1 Insights from Terrabacteria dataset

We investigated the use of this approach in prokaryotes by modeling different permutations of a Terrabacteria dataset to create scenarios that alter the number of taxa (and their identity) representing a given phylum while keeping other variables constant. Our results show two main trends: (i) the lower the number of taxa in a phylum the larger the variance in tree reconstruction and (ii) even with identical number of taxa, the identity of the species chosen to represent the phylum affects the reconstructed phylogeny. Among all species in our dataset, we identified a minimum of 15 (1 Deinococcus-Thermus, 5 Chloroflexi, and 9 Actinobacteria) that show consistently different phylogenies compared to the majority of the other species. Indeed, a re-analysis of our original dataset with only the species identified as problematic and without them produced two different topologies (Tree B and Tree A, respectively; Fig. 3a), suggesting that these species have unique genomic characteristics that are strong enough to alter the overall phylogeny. When we tested for possible phylogenetic artifacts driven by alignment length, compositional biases, and branch lengths we found no strong correlation with topological changes (Supplementary Data). However, multiple other factors have been proposed as potential sources of tree incongruences with permutated datasets and our results suggest that the species we identified as producing alternate trees are more likely to be sensitive to one or more of these sources of bias (Philippe *et al.*, 2011; Som, 2014).

In addition to these species, two Firmicutes (*Coprothermobacter proteolyticus* (CTP) and *Thermodesulfobium narugense* (TSN)) were consistently phylogenetically displaced, as they did not cluster within their phylum in the majority of the trees produced. This result supports previous findings based on large TOL analyses (Lang *et al.*, 2013) and more specific phylum-level evolutionary characters (Zhang and Lu, 2015). For example, Lang *et al.* (2013) show differences in their placement based on reconstructed methods: in their 16S rRNA trees, these species group together and are part of a basal polytomy of Bacteria, while in their ML trees TSN is sister to Dictyoglomus and those two are sister to a clade of Thermotogae and CTP. Another study that focused on these species in particular showed an indel pattern unique to them, questioning their classification with the Firmicutes (Kunisawa, 2015). Given the mounting evidence of incongruences of CTP and TSN with the remaining Firmicutes, we agree with previous studies that suggested that their classification should be revised (Pavan *et al.*, 2018).

Although the trends we identified are generally applicable to all scenarios we tested, a few exceptions exist. For example, Chloroflexi has lower discord among permutations than Deinococcus-Thermus and Actinobacteria and the variance within the discord is lower (fig 7, panels l and n). This is in agreement with the identification of molecular signatures within the Chlorofexi that show 28 conserved signature indels (CSIs) specific to different clades of Chloroflexi (Gupta, 2010). The evolutionary relationships between these CSIs indicate that the 4 different genera analyzed here form two clusters: Choroflexus/Roseiflexus and Dehalococcoides/Dehalogenimonas (Gupta *et al.*, 2013). While similarity between the members of each cluster is high, comparisons between clusters show lower similarity as seen also in our MID sampling permutations (Fig. 7: panel e). Our results also show that our large group permutations produced greater discord than permutations of our small groups for both HIGH and LOW sampling scenarios (Fig. 7: purple violin in panels j-l). This suggests that the presence of large number of taxa does not shield the group from phylogenetic instability when the group size is reduced by a few taxa (10%) (Fig. 7: panel j).

### 4.2 Extension to larger datasets

Based on these results obtained with a phylogeny of five phyla, we applied our procedure to one recent TOL composed of > 120 phyla (Hug *et al.*, 2016). In this case, our prediction was that we would observe less phylogenetic instability because the effect of permutating a single phylum (Deinococcus-Thermus, 6 species) would be diluted by the presence of many other groups. Instead, we found that the amount of destabilization of the phylogeny was ten-fold higher than in our smaller example. Thus, the effect of contrasting signals in a single, small phylum has far-reaching effects across a phylogeny.

Overall, these results point to the importance of testing phylogenetic stability even when using very large datasets. In the absence of such analyses, the phylogeny produced is driven by the signal present in the majority of the species, whether accurate or not, while alternate signals remain hidden. These alternative phylogenies could, however, provide important information on the evolutionary histories of specific lineages and genes and identifying the species that produce them is a necessary first step.

### 4.3 Implications

Despite significant advancements in computation and the feasibility of working with large genomic data, phylogenetic reconstruction efforts are unable to resolve the backbone phylogeny of many extant clades. Thus, a straightforward approach to test phylogenetic robustness under multiple scenarios is imperative, especially in very large datasets that require large computational times to analyze and are still sensitive to dataset changes. Although our study makes no claims on the ‘true’ phylogeny of the Terrabacteria superphylum, we showed that simply using a relative framework it is possible to determine the robustness of a dataset to taxon sampling and identify species that show conflicting phylogenetic signals. In particular, we have shown that it is important to iterate through many possible combinations of the same number of species to disentangle signals produced purely by the number of taxa from those produced by unique properties in each species genome or evolutionary history. These species can then become the target of further analyses to evaluate the presence of unique genomic properties, such as unique compositional biases or different gene evolution histories. This is especially important for clades represented by only a few genomes as any bias within these genomes will have a stronger overall weight on the reconstructed phylogeny of even distantly related groups. The presence of sparsely represented clades in prokaryotes is relatively common and is likely to remain so for the foreseeable future considering that over 70% of the sequenced and on-going sequencing projects represent only three phyla (Actinobacteria, Firmicutes, and Proteobacteria) (NCBI, 2017). Thus, many other phyla will remain represented at a much lower level and more susceptible to taxon sampling issues similar to those identified here.

